# LRX Proteins play a crucial role in pollen grain and pollen tube cell wall development

**DOI:** 10.1101/223008

**Authors:** Tohnyui Ndinyanka Fabrice, Hannes Vogler, Christian Draeger, Gautam Munglani, Shibu Gupta, Aline G. Herger, Paul Knox, Ueli Grossniklaus, Christoph Ringli

## Abstract

Leucine-rich repeat extensins (LRXs) are chimeric proteins containing an N-terminal leucine-rich repeat (LRR) and a C-terminal extensin domain. LRXs are involved in cell wall formation in vegetative tissues and required for plant growth. However, the nature of their role in these cellular processes remains to be elucidated. Here, we used a combination of molecular techniques, light microscopy, and transmission electron microscopy to characterize mutants of pollen-expressed *LRXs* in *Arabidopsis thaliana*. Mutations in multiple pollen-expressed *lrx* genes causes severe defects in pollen germination and pollen tube (PT) growth, resulting in a reduced seed set. Physiological experiments demonstrate that manipulating Ca^2+^ availability partially suppresses the PT growth defects, suggesting that LRX proteins influence Ca^2+^-related processes. Furthermore, we show that LRX protein localizes to the cell wall, and its LRR-domain (which likely mediates protein-protein interactions) is associated with the plasma membrane. Mechanical analyses by cellular force microscopy and finite element method-based modelling revealed significant changes in the material properties of the cell wall and the fine-tuning of cellular biophysical parameters in the mutants compared to the wild type. The results indicate that LRX proteins might play a role in cell wall-plasma membrane communication, influencing cell wall formation and cellular mechanics.

## Introduction

Upon germination of the pollen grain (PG), the pollen tube (PT) grows by the highly coordinated apical addition of newly synthesized cell wall materials and apical cell wall expansion driven by turgor pressure. The PT is one of the best models to study plant cell biology. The fine-tuned deposition of plasma membrane and cell wall components, and the spatiotemporally coordinated establishment of interactions between them, is crucial for shape generation (Geitmann, 2010) and sustained PT growth (McKenna et al., 2009). A crucial player in PG germination and PT growth is Ca^2+^, which regulates the dynamics of many cellular events including exo/endocytosis (Steinhorst and Kudla, 2013) and cell wall rigidity (Hepler et al., 2013).

The growth of plant cells depends on the delicate coordination between extracellular events occurring in the cell wall and intracellular cytoplasmic responses. This requires that plant cells can sense and integrate changes in the cell wall and relay them to the cytoplasm, a role typically played by transmembrane proteins with extracellular and cytoplasmic domains. Such proteins can interact with constituents of the cell wall to modulate their activity and/or convey signals into the cell (Ringli, 2010a; Wolf et al., 2012). For instance, wall-associated kinases (WAKs) bind to pectins in the cell wall and regulate osmotic pressure (Kohorn et al., 2006; Brutus et al., 2010). The receptorlike kinase THESEUS1 monitors changes in the cell wall caused by a reduced cellulose content and induces secondary changes such as lignin deposition (Hématy et al., 2007). Some LRR-receptor proteins, such as FEI1 and FEI2, influence cell wall function and cellular growth properties by affecting cell wall composition (Xu et al., 2008). The further identification and characterization of extracellular components that interact with and relay information to membrane partners will serve to elucidate the complex network of signal integration and transduction events that coordinate plant cell growth and morphogenesis.

Genome analyses in Arabidopsis has identified an eleven-membered family of leucine-rich repeat extensin (*LRX*) genes specialized into two phylogenetic clades: four “reproductive” (*LRX8-11* also known as *AtPEX1–4*, expressed in pollen) and seven “vegetative” (*LRX1-7*, expressed in vegetative tissue) *LRX* genes (Baumberger et al., 2003a). Henceforth, we use the gene symbols *LRX8-LRX11* to avoid confusion with Arabidopsis peroxin (*AtPEX*) genes involved in peroxisome biogenesis (Distel et al., 1996). LRXs are proteins containing a signal peptide, an N-terminal (NT) domain preceding a leucine-rich repeat (LRR) domain, which is joined to a C-terminal extensin (EXT) domain by a cysteine-rich motif (Figure 1A, Figure S1A). For simplicity, the region from the start of the N-terminal domain to the end of the cysteine-rich region is called the LRR as previously defined (Baumberger et al., 2001). The LRR domain is thought to bind an interaction partner, while the extensin domain, which has the typical features of the extensin-class of structural hydroxyproline-rich glycoproteins (HRGPs) (Baumberger et al., 2003a), anchors the protein in the cell wall (Baumberger et al., 2001; Ringli, 2010b). LRX8-LRX11 share a high similarity in the LRR domain, whereas the extensin domains are quite diverse (Figure S2). While the function of the LRR domain is strongly sequence-dependent, previous analyses have shown that the repetitive nature of the extensin domain is important rather than the exact sequence *per se* (Baumberger et al., 2003b; Ringli, 2010b; Draeger et al., 2015). LRX proteins have been shown to modulate lateral root development (Lewis et al., 2013), cell wall assembly, and cell growth in different tissues (Baumberger et al., 2001; Baumberger et al., 2003b; Draeger et al., 2015) which, based on their structure, was suggested to involve a regulatory and/or signalling function (Ringli, 2005). However, the nature of the interaction and the candidate regulatory and/or signalling processes that involve LRX proteins remain unknown.

**Figure 1.**
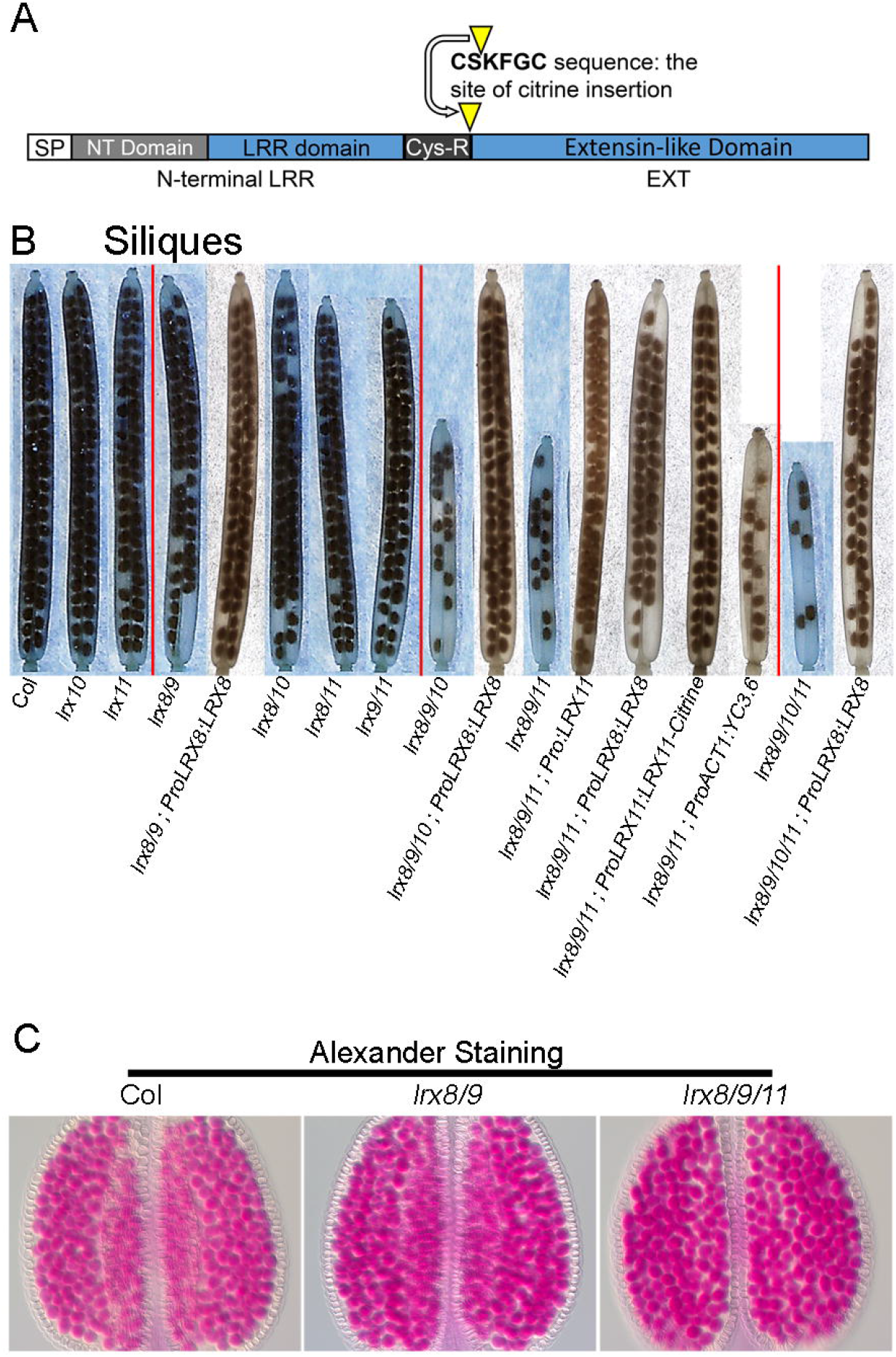
Seed set and pollen viability. **(A)** Schematic representation of the LRX proteins indicating the site of Citrine insertion. **(B)** Representative images of fully developed siliques of the wild type and different single, double, triple, quadruple *lrx* mutants, and complemented lines. Most severe defects are observed in the triple and quadruple mutants. Seed set is partially or fully restored in complemented lines (mutant background and gene used for complementation are separated by a double colon). **(C)** Alexander staining of anthers showing comparable pollen viability in the wild type and *lrx8/9* and *lrx8/9/11* mutants.

To address these issues and the relevance of pollen-expressed *LRXs* for plant reproduction, we isolated and characterized *lrx* mutants and found them to show reproductive defects, such as male sterility and reduced seed set. Our results reveal that LRXs are important players in the regulation of cell wall composition, structure, and mechanical properties. Vesicle dynamics and integration of new cell wall materials are affected in the *lrx* mutants, and these phenotypes can be suppressed by modulating Ca^2+^ availability. Based on these results, we propose a role of LRX proteins in PG and PT formation by influencing structures in and mechanical properties of the cell wall. The indications of changes in Ca^2+^-related aspects and vesicle dynamics, together with the observed association of the LRR domain of LRXs with the plasma membrane, indicate that these proteins might function as signalling components that link processes in the cell wall and the plasma membrane.

## Results

### *LRX8, LRX9, LRX10*, and *LRX11* are redundantly required for pollen tube function

We selected T-DNA insertion lines from the SALK library (Alonso et al., 2003) with insertions in *LRX8* (At3g19020; SALK_001367), *LRX9* (At1g49490; SALK_136073), *LRX10* (At2g15880; SALK_087083), and *LRX11* (At4g33970: SALK_030664), i.e. all the *LRX* genes expressed in pollen (male gametophyte). The T-DNA insertions interrupt the LRR-domain coding sequence (Figure S1A). Different combinations of double, triple, and quadruple mutants were produced through crosses and genotyping. Quantitative RT-PCR was used to test for abundance of the mRNAs in wild type and *lrx* quadruple mutants, using primers corresponding to regions 5’ upstream and 3’ downstream of the T-DNA insertion sites. Amplification of sequences 5’ and 3’ of the insertion sites was reduced in the *lrx* mutants by 30%-80% and over 95%, respectively (Figure S1B). Considering the importance of the LRR domain for LRX function (Baumberger et al., 2001; Ringli, 2010b), truncated LRX proteins with only parts of the LRR domain are very likely to be non-functional. This assumption is supported by the recessive nature of the *lrx8-11* mutations observed in the analyses described below. Reduced seed set was observed in various mutant combinations (Figure 1B, Figure S1C). Functional redundancy and synergism between the *LRX* genes was revealed when multiple mutations were combined. Reduced seed set was most severe in the quadruple mutant. Complementation of double, triple, and quadruple mutants with the genomic copies of *LRX8* or *LRX11* largely restored seed set (Figure 1B). Attempts to clone *LRX9* and *LRX10* failed, which excluded testing complementation with these two genes. Reciprocal crosses revealed 100% transmission of the mutations through the female, but not the male gametophyte (Table S1), indicative of a defect in PG/PT development or function. Alexander staining of mature PGs revealed that pollen viability in all the *lrx* mutants was comparable to the wild type (Figure 1C). Hence, the reduction in male transmission efficiency due to mutations in the *LRX* genes can be ascribed to a post maturation event, such as PG germination, PT growth, PT guidance, PT reception, and/or double fertilization. Given the redundancies observed between different combinations of the *lrx* mutants, and very poor germination of the quadruple mutant, we considered a subset (*lrx11, lrx8/9, lrx8/9/10*, and *lrx8/9/11*) for most of the characterization described below.

### LRX proteins regulate pollen germination and pollen tube growth

When germinated *in vitro* for 2 h, the *lrx* mutant pollen showed varying germination rates (Figure 2A). Instead of germinating, mutant PGs frequently burst, with up to 70% of *lrx8/9/10/11* grains bursting after 5 h *in vitro* (Figures S3A and S3B). Early time points were used for these analyses to have a developmental stage comparable to later experiments. The regulated uptake of water and swelling is required for PG germination and PT growth (Sommer et al., 2008). During imbibition in pollen germination medium (PGM), *lrx8/9/11* PGs swelled and burst (Movie S1). Also *in vivo*, ultrastructural analysis of germinating pollen showed aberrant structures in mutant PGs and PTs on stigmatic papillae such as cytoplasm remaining in the mutant PGs while in wild-type PGs, cytoplasm is transported along the PT (Krichevsky et al., 2007). Very few PTs eventually grew through the papillar apoplast into the ovary (Figure 2B, Figure S3C), thus accounting for the reduced seed set.

**Figure 2.**
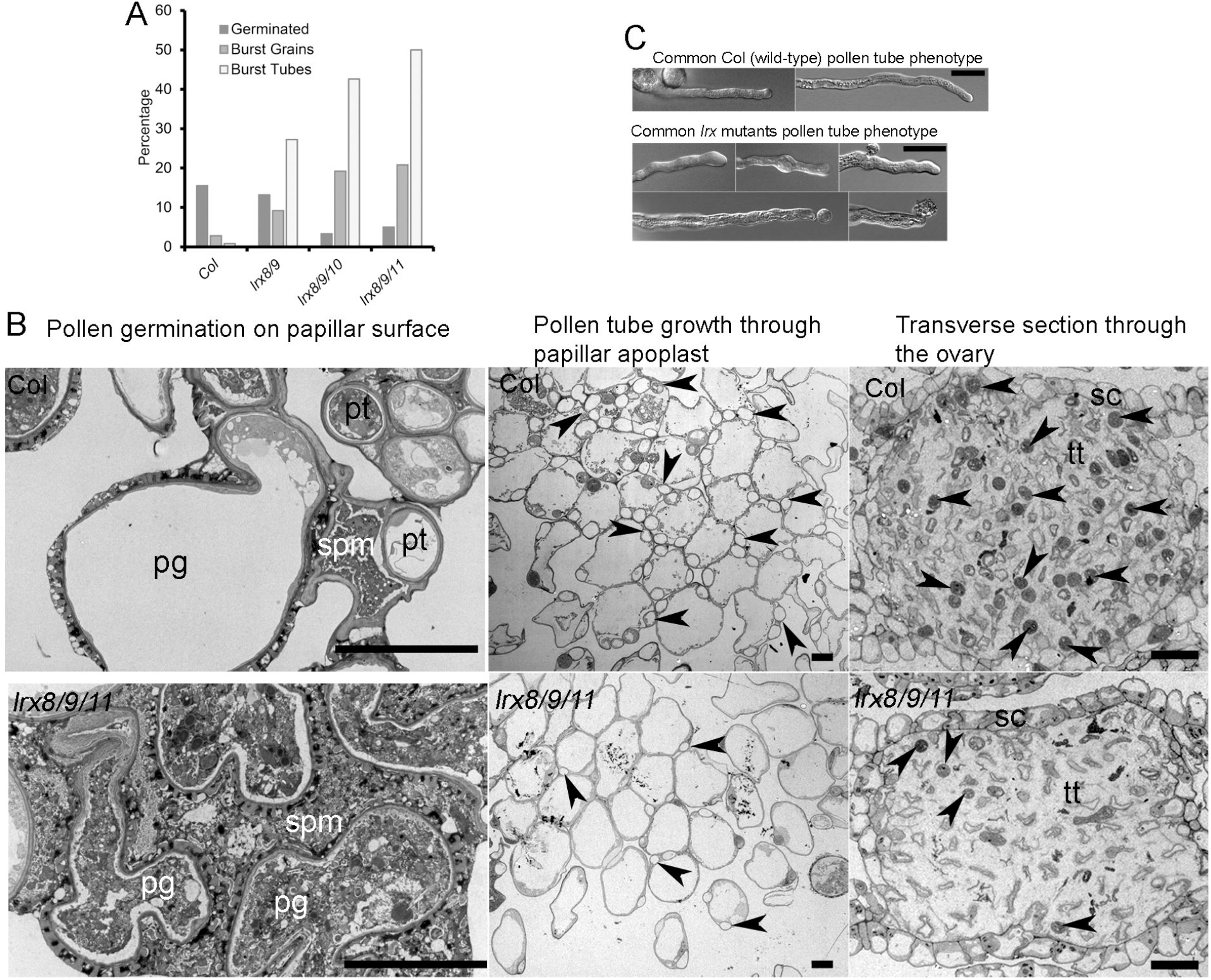
Pollen germination and pollen tube growth. **(A)** Percentage germinated PGs, burst grains, and burst tubes after 2 h of incubation *in vitro*, showing higher proportion of burst PGs and PTs in the *lrx* mutants. Percentage of burst PTs is based on the germinated PTs. Non-germinated PGs were distinguished from germinated PGs by the absence of a detectable PT. **(B)** Transmission electron micrographs of transverse sections through stigma/papilla surface, stigmatic papillae, and transmitting tract of the ovary. While wild-type PGs germinate and PTs grow, *lrx8/9/11* mutant grains mostly burst, discharge their content into the stigmatic papillar matrix (spm), and shrink. Eventually, compared to the wild type, fewer *lrx8/9/11* mutant PTs (some indicated by arrows) grow through the papillar apoplast into the ovary transmitting tract (tt) that is surrounded by septum cells (sc). **(C)** Typical wild-type PTs (mostly regular cylindrical) and *lrx* PTs (bulging, bursting, budding) phenotypes are shown. pg=PG, spm=stigmatic papillar matrix, tt= transmitting tract, sc= septum cells. scale bar B = 10 μm, C = 20 μm.

The bursting of *lrx* PGs prompted us to examine whether there were any visible ultrastructural alterations in mature PGs. While the organization of subcellular structures in the cytoplasm of mature PGs looked similar in mutants and the wild type, the intine wall in *lrx* mutants was split into two by a band of electron-dense material, which was absent in the wild type (Figure S4). This data fits the previously proposed role for an LRX-type protein in the maize pollen intine (Rubinstein et al., 1995). Thus, the impaired germination of *lrx* mutant PGs indicates that LRX proteins are involved in the fine-tuning of the rapid and dynamic cellular processes required for successful PG germination by regulating the formation of the intine wall.

In PTs, *lrx* mutants showed varying frequencies of diverse phenotypes, mainly swelling at intervals, vesicle budding at the apex, and PT bursting (Figures 2A and 2C). Mutant PTs exhibited an intermittent growth behaviour, i.e. phases of rapid growth interspaced by phases of very slow or no growth (growing PTs would stop growth for varying periods of up to 30 min before resuming growth). Time-lapse imaging of PTs stained with the lipophilic styryl dye FM1-43 (Betz et al., 1992; Vida and Emr, 1995) revealed release of cellular contents (Movie S2) or budding of vesicles in the apical region (Movie S3) in *lrx8/9/11* PTs. Consequently, the PT growth rate was significantly reduced in the *lrx* mutants compared to the wild type, where PT growth was continuous as illustrated in kymographs that monitor progression of the PT tip over time (Figure 3C). The apical accumulation of secretory vesicle content has been characterized during exocytic discharge/integration of new cell wall material, and precedes and determines the peak in PT growth for which it is required (McKenna et al., 2009). The aberrant discharge of vesicles in the *lrx* mutants are indicative for a role of LRX proteins in the coordinated integration of new cell wall material during PT growth.

**Figure 3.**
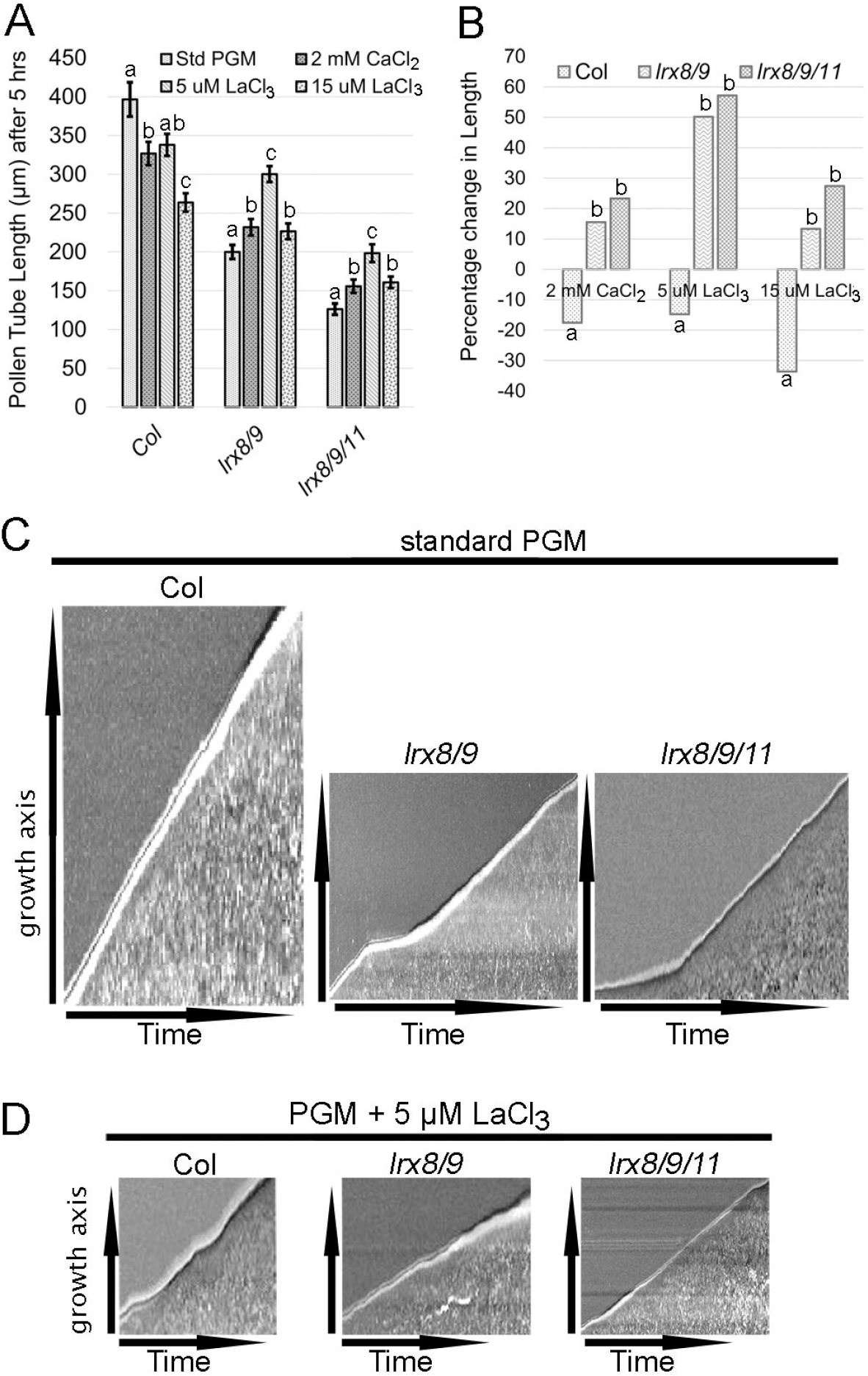
Pollen tube growth in different Ca^2+^ regimes. **(A)** Average length of PTs in standard (std) PGM (containing 5 mM CaCl_2_), PGM containing reduced [Ca^2+^] (2 mM CaCl_2_), and in PGM supplemented with 5 μM and 15 μM LaCl_3_. While lower [Ca^2+^] or LaCl_3_ treatments reduced PT growth in the wild type, these regimes improved PT growth in *lrx8/9* and *lrx8/9/11* mutants (n≥200; error bar = s.e.m.; different letters indicate significant differences, *t*-test, P<0.05). **(B)** The percentage change in PT length under different Ca^2+^ regimes compared to their corresponding values in standard PGM. The highest positive effect is obtained with the *lrx8/9/11* mutant at 5 μM LaCl_3_. **(C)** Kymographs showing continuous and intermittent growth of wild-type and *lrx* PTs, respectively, in standard PGM. **(D)** In PGM containing 5 μM LaCl_3_, continuous growth is slightly perturbed in the wild type, while intermittent growth is partially restored to continuous growth in *lrx* mutants.

### LRX proteins modulate the composition and ultrastructure of the pollen tube cell wall

Given the observed impairment of the integration of new cell wall material in *lrx* PTs, we used a panel of monoclonal antibodies (mAB) for immunofluorescence studies of some major ER/Golgi- and plasma membrane-synthesized cell wall components in wild-type and *lrx* mutant PTs. We used, for the ER/Golgi-synthesized wall epitopes, JIM20 against extensins (Smallwood et al., 1994), LM2 against arabinogalactan-proteins (AGPs) (Yates et al., 1996), LM6 against the arabinan domain of rhamnogalacturonan I (RG-I) (Willats et al., 1998), LM19 and LM20 against unesterified and methyl-esterified homogalacturonan, respectively (Verhertbruggen et al., 2009), LM15 against xyloglucan (Marcus et al., 2008), and aniline blue to stain plasma membrane-synthesized callose. Interestingly, all the ER/Golgi-synthesized cell wall components were reduced in the *lrx8/9, lrx8/9/11*, and *lrx8/9/10* mutants compared to the wild type (Figure 4A, Figure S5). A frequent feature in *lrx* mutants was a locally increased signal of some wall epitopes associated with bulged regions of the PT, a likely reflection of the intermittent growth behaviour. In addition, the released cytoplasmic contents were usually heavily labelled for cell wall epitopes with different antibodies, indicating the discharge of wall materials (Figure 4B). This overall reduction of wall epitopes may be due to a net reduction in the rate of synthesis of cell wall components, the masking of the epitopes by altered bonding patterns and reducing the spaces between cell wall polymers, or a cytoplasmic accumulation and/or failure in the integration of exocytic cell wall components into the cell wall (see below). When PTs were treated with xyloglucanase to digest xyloglucan and facilitate diffusion of mAbs into the cell wall, the labelling of cell wall epitopes was still lower in *lrx* mutants than the wild type (Figure 4A). Frequently, labelling of wall epitopes that were still within the cytoplasm was observed in the mutants. These data suggest that the antibodies could penetrate the cell wall and, hence, the masking of cell wall epitopes by reduced spacing was likely not responsible for the decreased labelling seen in *lrx* mutants. The *lrx8/9* and *lrx8/9/11* mutants accumulated plasma membrane-synthesized callose while aniline blue staining was weaker in the wild type. Frequently, the labelling for callose in the mutants spanned the entire cell wall of the PT, contrary to the wild type where labelling was restricted to the shank (Figure 4A) as previously described (Dardelle et al., 2010; Chebli et al., 2012).

**Figure 4.**
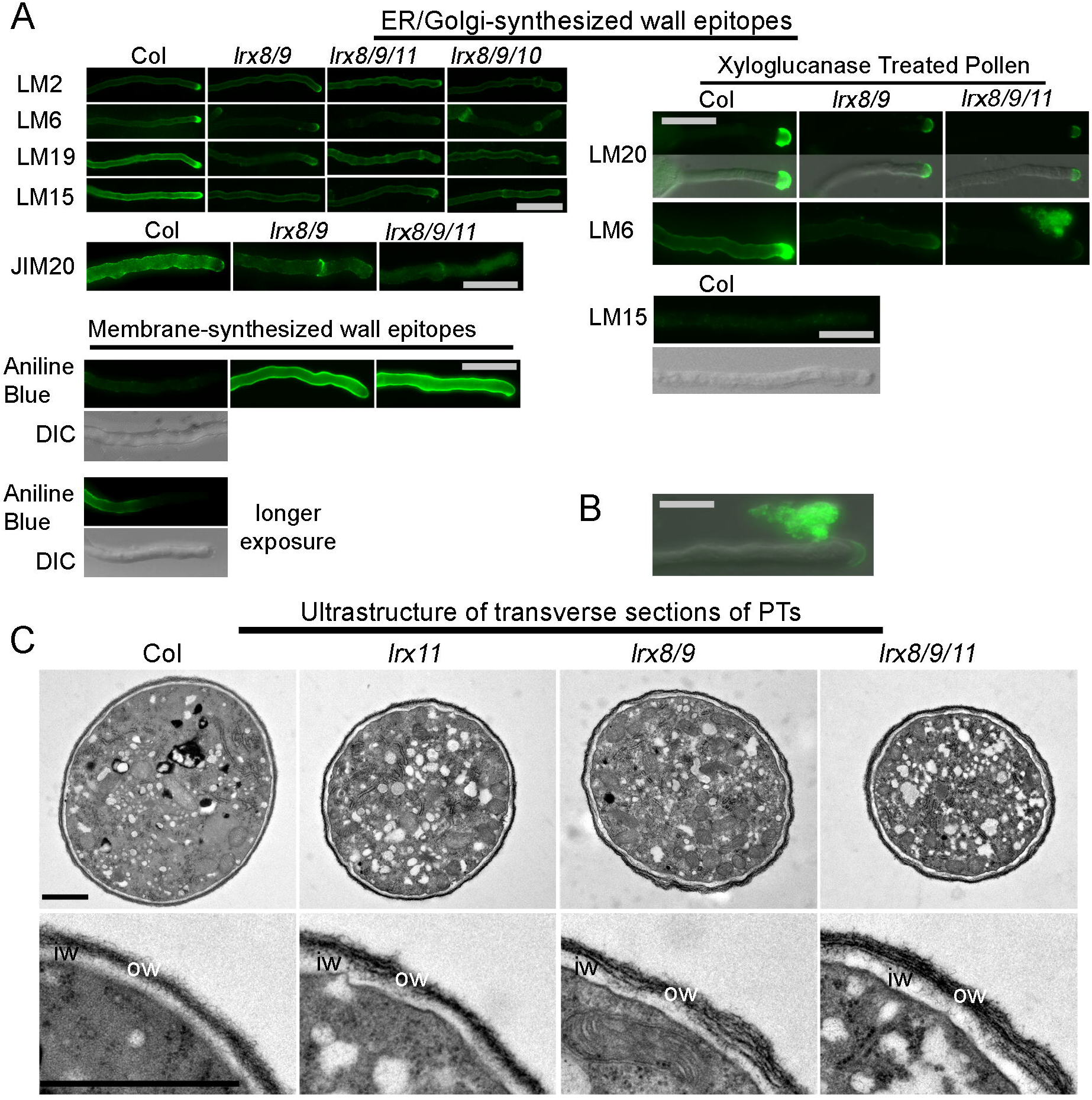
Immunolabelling and ultrastructural analyses of pollen tube cell walls. **(A)** The labelling of ER/Golgi-synthesized cell wall components (with LM2, LM6 LM19, LM15, and JIM20) and plasma membrane synthesized cell wall components (with aniline blue). The labelling for ER/Golgi-synthesized wall components is significantly weaker in *lrx8/9, lrx8/9/11*, and *lrx8/9/10* compared to the wild type. The labelling for plasma membrane-synthesized callose (aniline blue) in mutants is stronger than in the wild type and also found at the tip. Even longer exposure of wild-type PTs demonstrates callose labelling in the shank but not at the tip. Xyloglucanase treated PTs still show significantly lower labelling for pectin (LM20, LM6) compared to the wild type, and no labelling for xyloglucan (LM15). DIC captures are shown for aniline-stained wild-type PT and LM15-labelled PT treated by xyloglucanase to show position of the PTs. **(B)** Cytoplasmic content released from the PT strongly stain for cell wall components, as shown here for pectin (LM6). **(C)** TEM transverse sections of PTs. The mutant PT outer cell walls are more loose/fibrous and the inner wall thicker, reflecting the higher accumulation of callose (strongest in *lrx8/9/11)* compared to the wild type. Scale bar A = 20 μm, B = 10 μm, C = 1 μm

Given these changes in the cell wall structure of *lrx* mutants, we analyzed the ultrastructure by transmission electron microscopy (TEM). Transverse sections of PTs revealed ultrastructural changes in the mutants including a loosely packed, fibrous outer wall and a thicker, electron-weak, callosic inner wall, features that were most conspicuous in the *lrx8/9/11* triple mutant (Figure 4C). Thus, the observed altered representation of wall epitopes in the *lrx* mutant is associated with a modified ultrastructure of the PT cell wall. The more fibrous wall is likely a consequence of less ER/Golgi-produced cell wall matrix material being deposited.

### The *lrx* mutants show altered vesicle dynamics

We investigated whether the reduced abundance of ER/Golgi-synthesized cell wall components could be due to a defect in the transport or cytoplasmic accumulation of secretory vesicles. First, the wild type and *lrx8/9/11* triple mutants were transformed with a *pLAT52::GFP-fABD2* construct to visualize actin filament organization, which is important for the movement of secretory vesicles to the apical growth region, tip growth, and wall organization (Zhang et al., 2010), a prerequisite for sustained PT growth. We observed a similar actin cytoskeleton orientation in the shank of wild-type and mutant PTs (Figure S6A), suggesting that the trafficking of ER/Golgi-synthesized wall components to the apex was likely unaffected in the *lrx* mutants. Next, we used an established method to investigate exocytosis efficiency (Samalova et al., 2006). Wild-type and *lrx8/9/11* plants were stably transformed with a *pACT1::nlsRm-2A-secGf* construct. The 2A peptide sequence of nlsR_m_-2A-secG_f_ allows the cleavage of the polyprotein into equimolar amounts of the secGFP and nlsRFP protein moieties. The nlsRFP moiety accumulates in the nucleus where it emits RFP fluorescence, while the secGFP moiety is exported to the cell wall, where its GFP fluorescence is poor due to the acidic pH (Samalova et al., 2006). A defective export process results in increased intracellular GFP and, hence, a lower RFP:GFP ratio. The ratio of RFP:GFP fluorescence is not significantly altered between mutant and wild-type PTs (Figures S6B and S6C). Thus, the apparent reduction of ER/Golgi-synthesized cell wall epitopes in the *lrx* mutants is not due to an accumulation of secretory vesicles in the cytoplasm, but possibly to a failure in the later steps of exocytosis, notably vesicular discharge and correct integration of new cell wall material into the expanding cell wall. This interpretation is supported by the uncontrolled vesicle budding and discharge of cell wall material into the surrounding medium (Movies S1–S3).

With the aberrant discharge of cell wall material and the consequent slower PT growth, especially in the *lrx8/9/11* triple mutant, we envisioned an increased accumulation of excess membrane in the apical plasma membrane. This would require endocytic recycling (Battey et al., 1999), leading us to investigate the rate of endocytosis using the lipophilic styryl dyes FM4-64 and FM1-43 (Betz et al., 1992; Vida and Emr, 1995). Quantification of FM4-64 fluorescence in the apical cytoplasm revealed a significantly increased rate of membrane uptake, and hence increased endocytosis, in the *lrx8/9/11* triple mutant (Figures S7A and S7B). The rate of endocytosis was comparable between the *lrx8/9* double mutant and the wild type. A similar result was obtained using FM1-43. The increased endocytosis in the *lrx8/9/11* mutant is likely induced by the PT to counteract over-accumulation of plasma membrane material that is caused by the slower PT growth at a normal rate of exocytosis.

### Altering Ca^2+^availability alleviates the *lrx8/9/11* PT growth phenotypes

Since Ca^2+^ in PTs regulates vesicle fusion (Camacho and Malhó, 2003), PT bursting at the micropyle (Iwano et al., 2012; Ngo et al., 2014), and is hypothesized to promote endocytosis (Zonia and Munnik, 2009), we speculated that Ca^2+^ dynamics might be altered in *lrx* mutant PTs. To investigate this possibility, the influence of modulated Ca^2+^ availability on PT growth was investigated. When growing PTs on PGM containing reduced [Ca^2+^] (2 μM instead of 5 μM in standard PGM), *lrx* mutant PTs showed increased tube length at 5 hrs post-germination, whereas wild-type PTs were negatively affected. Similarly, inhibiting Ca^2+^ channels by supplementing the PGM with 5 μM or 15 μM LaCl3 (a Ca^2+^ channel blocker) had a positive effect on *lrx* PT growth but negatively affected the wild type (Figures 3A and 3B). The intermittent growth phenotype of the *lrx8/9/11* PT visible on kymographs was largely restored to continuous growth at 5 μM LaCl3, whereas wild-type PTs grew less steadily (Figure 3D). Finally, the increased rate of endocytosis observed in *lrx* mutant PTs grown in the presence of 5 μM LaCl3 was reduced to wild-type levels (Figure S7B). Together, the observed correlation between the reduction of Ca^2+^ levels and the alleviation of the PT growth phenotype in the *lrx* mutant indicates a possible role of LRX proteins in Ca^2+^-related processes. To investigate the [Ca^2+^] dynamics in PTs, wild-type and *lrx8/9/11* triple mutant plants were transformed with the ratiometric Ca^2+^ indicator protein yellow cameleon 3.60 (YC3.60) (Nagai et al., 2004). Growing PTs have an intracellular [Ca^2+^] gradient with a peak in the apical cytoplasm required for PT growth (Iwano et al., 2009). Ratiometric analysis revealed the presence of this increased [Ca^2+^] at the apex in wild-type as well as mutant PTs (Figure 5). [Ca^2+^] oscillations (Pierson et al., 1996) were also similar in wild-type and mutant PTs. However, a strong increase in [Ca^2+^] was frequently observed in *lrx* mutant PTs that culminated in the bursting of the PTs. Together, these data reveal that the observed suppression of the *lrx* mutant phenotypes by reducing availability or transport of Ca^2+^ might be explained by an alteration in Ca^2+^ distribution in the *lrx* mutant PTs.

**Figure 5.**
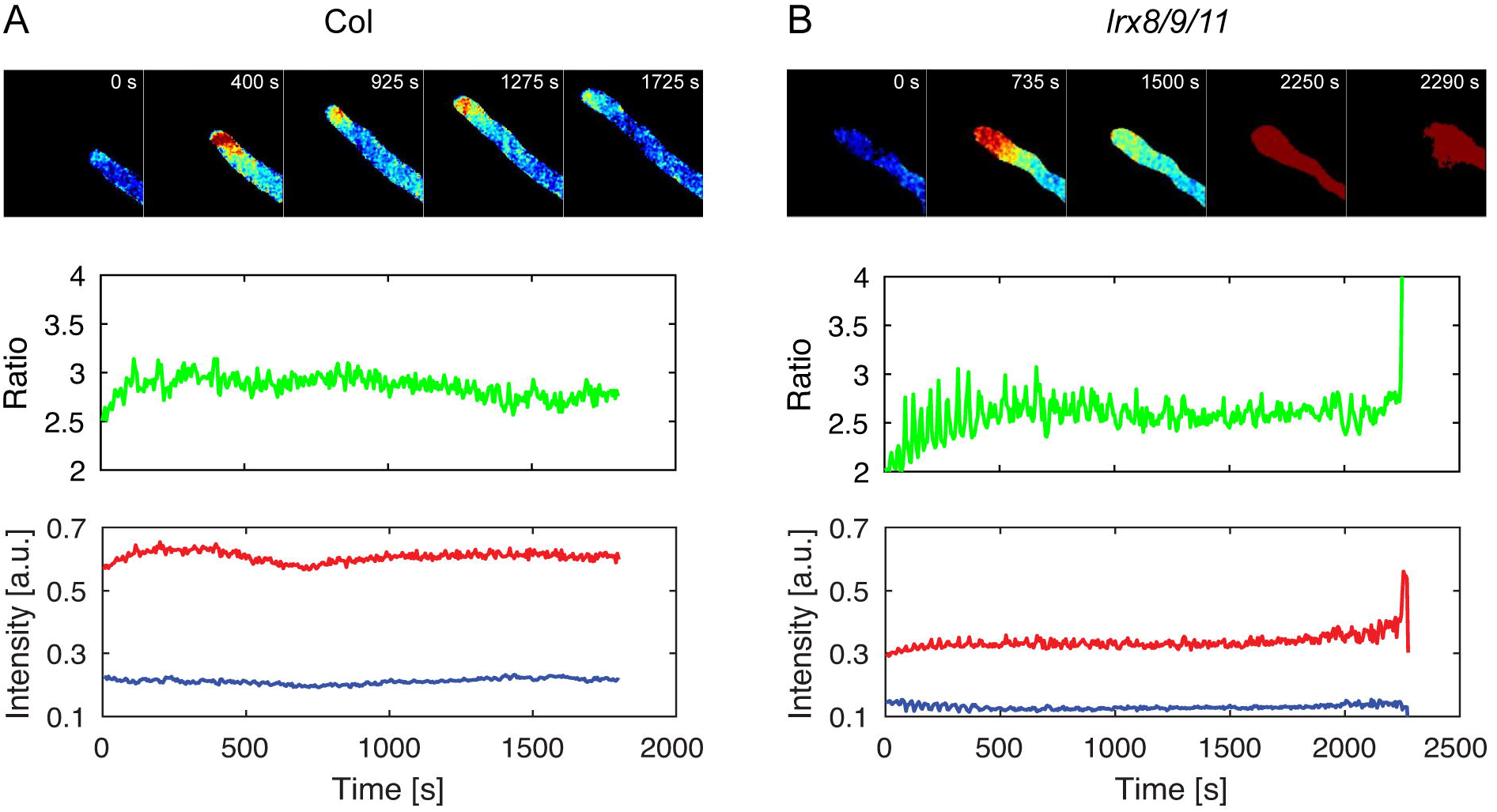
Visualization and quantification of intracellular Ca^2+^ dynamics. Time series of YC 3.60 fluorescence showing a [Ca^2+^] gradient with a tip-localized increase in wild-type **(A)** and *lrx8/9/11* **(B)** PTs (upper panel). A strong increase in [Ca^2+^] is seen in the mutant prior to bursting. Graphs show fluorescence of Ca^2+^-unbound YC 3.60 (blue line), Ca^2+^ -bound YC 3.60 (red line), and the ratio representing the Ca^2+^ signal (green line) with the spike in the mutant prior to bursting.

### The LRX N-terminal LRR-moiety associates with the plasma membrane

Given the ranges of cell wall/membrane-associated functions impaired in the *lrx* mutants, and the potential for protein-protein interaction of the LRR domain, we investigated its cellular localization. The coding sequence of the fluorescence protein Citrine was introduced near the C-terminal end of the cysteine-rich region (Figure 1A) of *LRX11* to produce *pLRX11::LRX11-Citrine*. Additionally, the extensin domain was deleted to produce *pLRX11::LRR11-Citrine*. The LRX11-Citrine but not the LRR11-Citrine restored seed set in the *lrx8/9/11* triple mutant back to double-mutant levels (Figure 1B, Figure S1C), indicating that the Citrine insertion does not obstruct the protein activity and that the extensin domain of LRX11 is required for protein function. Citrine fluorescence was observed in PGs and PTs, and a fraction of LRX11-Citrine remained in the PT cell wall after plasmolysis. By contrast, the LRR11-Citrine fusion protein was not observed in the cell wall after plasmolysis but appeared to retract with the plasma membrane (Figures 6A–6C). Investigating this observation further in PTs was technically not possible due to the low Citrine fluorescence in the transgenic lines. Therefore, a possible membrane association was further analyzed in lines expressing *LRR4-Citrine* under the *pLRX4* promoter (Draeger et al., 2015) that results in strong fluorescence in vegetative tissues. The N-terminal half of LRX4 shows high homology to LRX8-LRX11 (Figure S2) with 56-58% and 83-85% identical and similar positions, respectively. In these transgenic seedlings, Citrine fluorescence was associated with the plasma membrane after induction of plasmolysis (Figure 6D). To confirm this observation by an alternative approach, either total extracts or membrane fractions of wild-type and *pLRX4::LRR4-Citrine* transgenic seedlings were isolated and tested for the presence of LRR4-Citrine by Western blotting.

**Figure 6.**
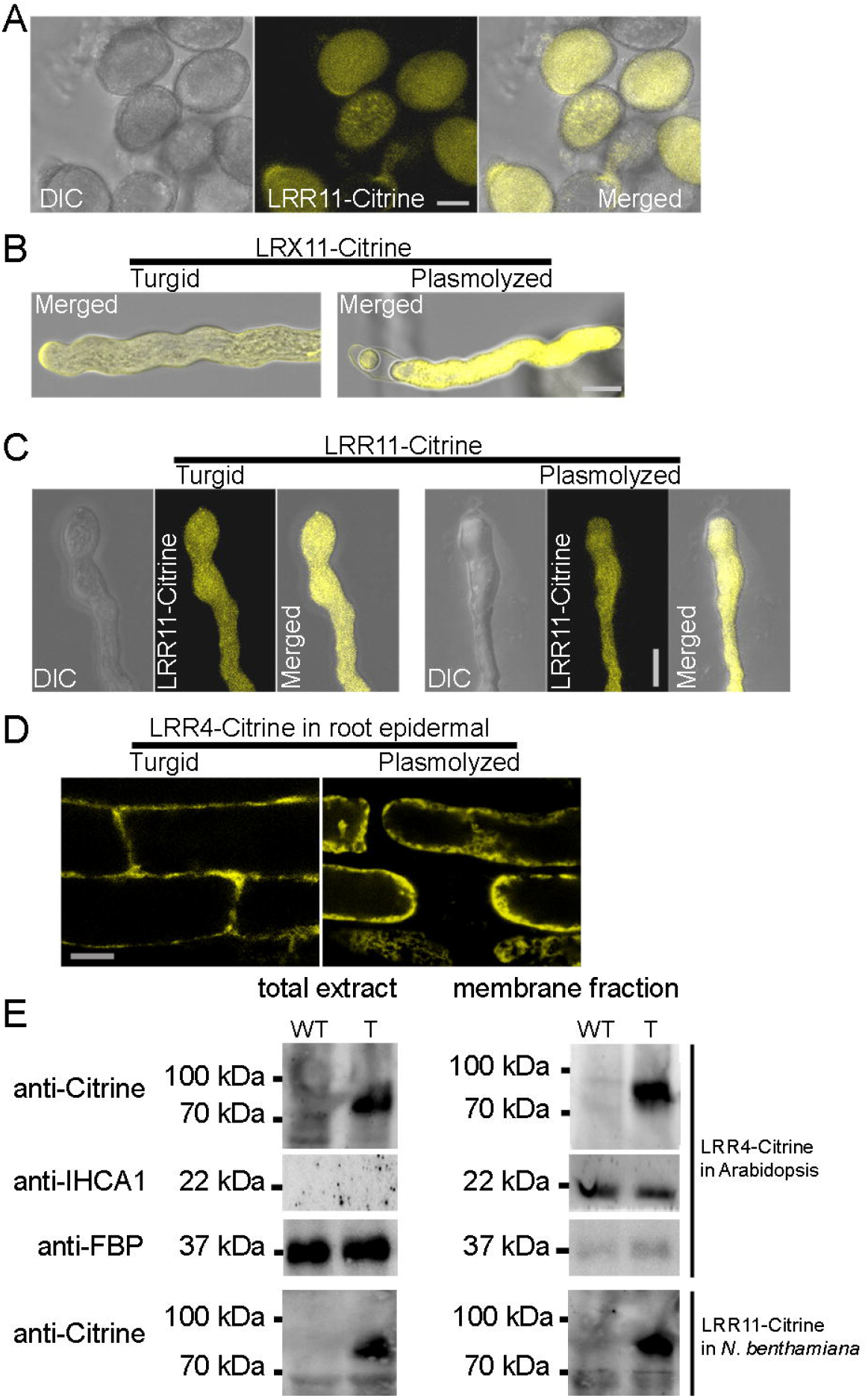
LRX-Citrine and LRR-Citrine localization. **(A)** LRR11-Citrine fluorescence in PGs. **(B)** LRX11-Citrine localization in cell wall and cytoplasm in turgid and plasmolyzed PTs. **(C)** LRR11-Citrine localizes to the cell wall-plasma membrane and cytoplasm in turgid PTs, but retracts with the plasma membrane and cytoplasm in the plasmolyzed as does **(D)** LRR4-Citrine in hypocotyl cells of *pLRX4::LRR4-Citrine* transgenic seedlings. Scale bar = 10 μm **(E)** Western blot of total extracts (left panel) and membrane fractions (right panel) of wild-type (WT) and *pLRX4::LRR4-Citrine* transgenic (T) seedlings probed with an anti-GFP, anti-LHC1a, or anti-FBP antibody to detect LRR4-Citrine, the membrane protein LHC1a, and the cytoplasmic protein FBP, respectively. Tobacco leaf material expressing LRR11-Citrine (T) and non-transgenic tobacco (WT) was purified in the same way and LRR11-Citrine also co-purified with the membrane fraction. Scale bar (A-D) = 10 μm

As shown in Figure 6E, LRR4-Citrine fusion protein was found in both fractions and migrated at the expected size of around 75 kDa, whereas no signal is observed in the non-transgenic control. The membrane-bound protein LHC1a (Klimmek et al., 2005) and the cytoplasmic protein FBP (Folate Binding Protein) showed opposing, strong enrichment in the membrane and total fraction, respectively, of both the transgenic and non-transgenic line, confirming successful preparation of the membrane fraction (Figure 6E). In parallel, total fractions and membrane fractions of tobacco plant material transfected with an *p35S::LRR11-Citrine* overexpression construct also revealed the LRR11-Citrine band, while no protein was detected in non-transfected material (Figure 6E). Together, these experiments suggests that the LRR domain of LRX proteins associates with the plasma membrane.

### The *lrx* mutations alter the fine-tuning of pollen tube mechanics

The turgor pressure and cell wall stiffness of PTs offer powerful explanatory principles to explain cellular growth. These principles are even more instructive when the mechanical characterization of cellular growth is interpreted in terms of the cell wall structure (Cosgrove, 2015) balancing the turgor pressure. We used the cellular force microscope (CFM) (Felekis et al., 2011; Vogler et al., 2013) in conjunction with a quasistatic continuum finite element model (FEM) to determine the biophysical properties of wild-type and *lrx* mutant PTs. The CFM measures the force required to create an indentation of a given depth into the PT. The apparent stiffness is the slope of the resulting force-indentation depth curve. CFM measurements taken at about 10 μm behind the PT tip showed a significant increase in the apparent stiffness of *lrx8/9* double and *lrx8/9/11* triple mutant PTs compared to the wild type (Figure S8). The FEM model converts the CFM output into the decoupled mechanical properties of turgor pressure and cell wall stiffness; where the latter is defined as the product of the Young’s modulus (Figure S8) and the cell wall thickness. The cell wall thickness determined by TEM showed a significant increase in the *lrx* mutants (wt = 151 ±22 nm, *lrx8/9* = 168±25 nm, and *lrx8/9/11* = 202±34 nm, mean±SD, P value ≤ 0.0001; Figure S8). The model revealed a significant increase in the mean turgor pressure in the *lrx8/9* (P value < 0.0001) and *lrx8/9/11* PTs (P value < 0.03), as well as a significant increase in the mean cell wall stiffness of *lrx8/9/11* PTs (P value < 0.0001) (Figure 7). Thus, it seems that mutations in the *LRX* genes significantly alter the biophysical properties of PTs. The range of turgor pressure and cell wall stiffness values between the 10^th^ and 90^th^ percentile followed the mean values in also showing a larger increase compared to the wild type. This large variation and/or higher mean turgor pressure and cell wall stiffness could indicate that the *lrx* mutant PTs are suboptimal structures while wild-type PTs are finely-tuned optimized structures. This is reinforced by our finding that *lrx* PTs exhibit a variety of abnormalities, growth rate reduction, intermittent growth, and a higher propensity to burst.

**Figure 7.**
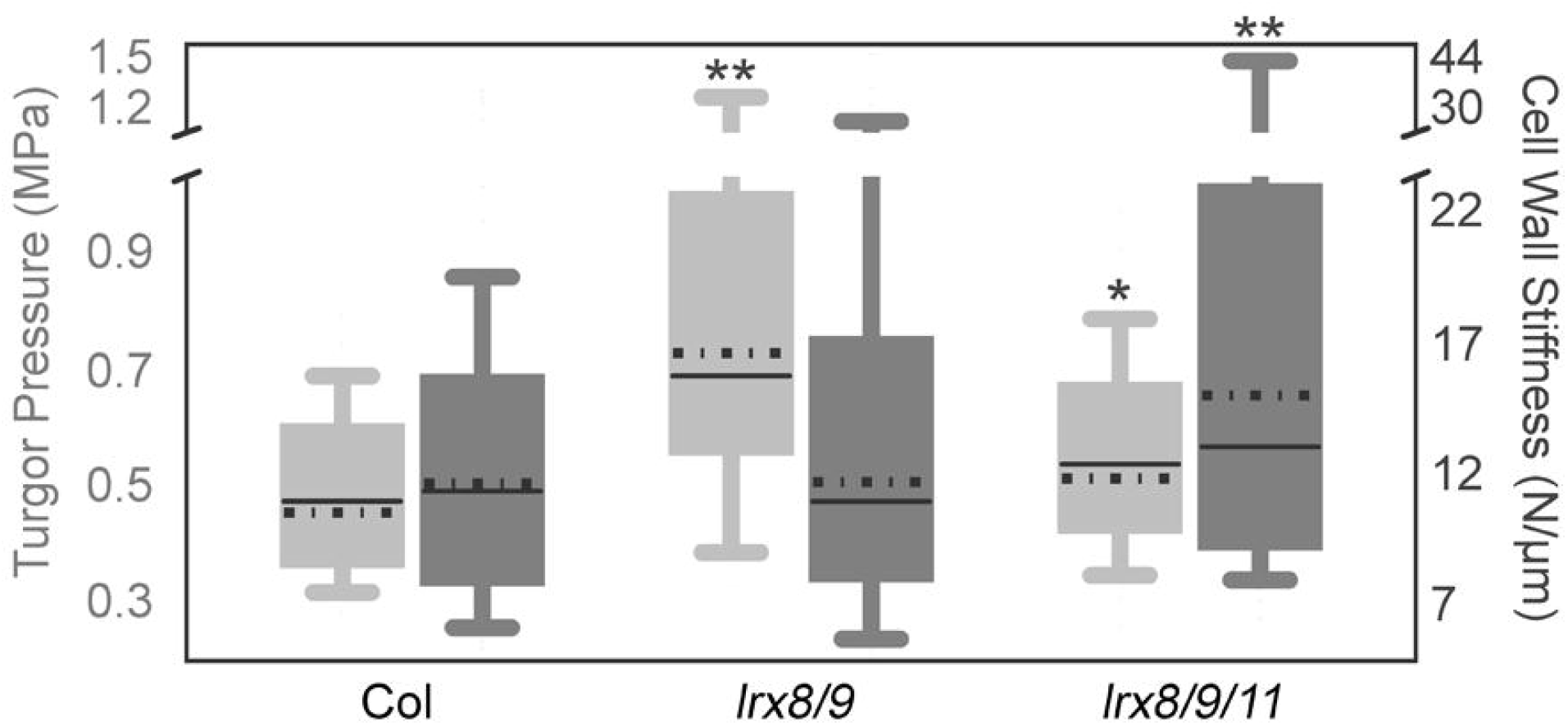
Biophysical properties of pollen tubes deduced by FEM-based modelling. Compared to the wild type, turgor pressure (light grey box) is significantly increased in the *lrx8/9* and *lrx8/9/11*. The stiffness of the cell wall (dark grey box) is significantly increased in the *lrx8/9/11* mutant compared to the wild type, which is similar in *lrx8/9*. Statistics: *t*-test, * = P < 0.03, ** = P < 0.0001, n≥50. In addition, the box plots show considerable skewness of turgor and cell wall stiffness in *lrx* mutants as revealed by the larger difference between the median (line) and the mean (stroked line) as well as the range of the whiskers compared to the wild type where the deviations are smaller. Given the skewness, the mean values shown are calculated from the log normalized data.

## Discussion

Cell growth in plants is coordinated through a plethora of processes in the cytoplasm, plasma membrane, and the cell wall, including the biosynthesis of cell wall materials, their transport to the plasma membrane, and their controlled deposition in the cell wall (Cosgrove, 2014). These events are tightly regulated both spatially and temporally, requiring signaling processes between cell wall and cytoplasm that allow the exchange and integration of information (Baluška et al., 2003). LRX proteins are extracellular players that are involved in these cellular processes, influencing endocytic membrane recycling, the release at the plasma membrane and integration of cell wall material into the existing cell wall and, consequently, cell wall properties and functions. We observed a reduction in ER/Golgi-originating cell wall matrix material in *lrx* mutant PTs as detected by immunostaining and a more fibrillar and less condensed outer PT cell wall structure as observed in TEM images. Although less likely, we cannot completely exclude that fundamental changes in many cell wall structures mask epitopes or inhibit the binding of the whole array of antibodies. As vesicle transport and exocytosis was not found to be altered in the mutants, it is most likely the subsequent, controlled deposition and coordinated integration of the newly synthesized wall components that seem affected in *lrx* mutants; a conclusion that is supported by the budding and discharge of vesicle contents observed in the apical region of *lrx* mutant PTs. LRX8-LRX11 are likely to contribute similarly to these processes, since combining different sets of *lrx* mutations have comparable effects.

Ca^2+^ regulates several cellular processes including the control of cellular growth. Its influence on vesicle dynamics (Picton and Steer, 1983; Camacho and Malhó, 2003; Steinhorst and Kudla, 2013), and the increased rate of endocytosis observed in the *lrx* mutants, led us to investigate the relevance of Ca^2+^-related processes for the *lrx* PT growth defects. The alleviation of the *lrx* mutant growth phenotypes by reducing Ca^2+^ uptake or by reduced [Ca^2+^] in the medium revealed a link between Ca^2+^ and the defects in *lrx* PTs. It remains an open question as to whether and how LRX proteins modulate these aspects of PT growth. Future experiments will determine whether LRX proteins influence Ca^2+^ distribution and possibly Ca^2+^ fluxes. Since a number of processes are affected, more detailed experiments are necessary to discern direct from secondary effects of the *lrx* mutations.

The full-length LRX proteins are insolubilized in the cell wall via the extensin domain that serves as an anchor (Baumberger et al., 2001; Ringli, 2010b). Interestingly, in the absence of the extensin domain, the N-terminal moiety shows association with the plasma membrane. Due to the low abundance of LRR11-Citrine in PTs, this analysis was done with LRR4-Citrine expressed in vegetative tissues, and with transiently expressed LRR11-Citrine in tobacco. The homology of the LRX proteins [(Baumberger et al., 2003a) and data shown] makes it likely that the different LRX proteins have the same functional principles, which is supported by the comparable properties of the two proteins. Membrane association is possibly mediated through interaction with a membrane-associated binding partner, which would establish LRX proteins as connectors of the cell wall with the plasma membrane. This hypothesis is supported by localization of an LRX-type protein of maize to the intine cell wall of PGs and the callose layer near the plasma membrane (Rubinstein et al., 1995). Linker activity has been attributed to several transmembrane proteins. For instance, WAKs interact with pectin in the cell wall (Brutus et al., 2010) and PERKs, based on the similarity of their extracellular domain to structural cell wall proteins, likely bind to cell wall components (Bai et al., 2009). LRXs, by contrast, would be a different type of linker protein since they have no transmembrane domain but are covalently connected to the cell wall (Baumberger et al., 2003a; Ringli, 2010). Whether LRX proteins interact directly with a plasma membrane-anchored component or indirectly via several other proteins remains an open question. The identification of the interaction partner(s) of LRX proteins will be essential for further elucidation of the membrane association, function, and mode of action of these proteins.

A defining effect of mutations in the *LRX* genes is a change in the cell wall structure, which can be quantified mechanically by material properties including cell wall stiffness, and consequently the rigidity of the entire PT as suggested by its apparent stiffness (Vogler et al., 2013). The FEM model predicts significantly higher turgor pressure in both *lrx8/9* and *lrx8/9/11* mutants, significantly higher cell wall stiffness in *lrx8/9/11*, and a large range of these mechanical properties in both *lrx* mutants compared to the wild type. This seems to provide a foundation for the lower growth rate and higher frequency of aberrant PT phenotypes. Furthermore, it should be noted that the experimentally measured input parameters (cell wall thickness, PT diameter, and apparent stiffness) and the output properties turgor pressure and cell wall stiffness showed a skewed distribution in the *lrx* mutants (large deviation between mean and median). This skewness can be attributed to the nontrivial percentage of *lrx* mutant PTs that burst before they could be used for CFM or TEM analyses, which should - in theory - reduce the number of extreme values instead of increasing it. Therefore, apparently only the best performing *lrx* mutants were measured, but even these seem suboptimal compared to the wild type. The variability of PT growth in *lrx* mutants likely reflects the ability of the PTs to sense defects in their cell wall structure/function and induce compensatory changes, as does THESEUS1 in sensing cellulose deficiency in *procuste* mutants (Hématy et al., 2007). In *lrx* mutants, the increased deposition of callose, even at the apex, is possibly a compensation for the PTs to overcome deficiencies in cell wall structure and stabilize the cell wall (Parre and Geitmann, 2005).

In conclusion, these analyses demonstrate that *LRX* genes have an important function in male gametophyte development and reveal possible processes that are influenced by the LRX proteins. This work confirms previous findings that these proteins are important for cell wall development (Baumberger et al., 2001; Draeger et al., 2015). The mechanism underlying the observed defect in the cell wall seems to involve fundamental cellular events that occur at the plasma membrane/cell wall interface. We suggest that LRX-type proteins serve as candidates for linking the plasma membrane and the extracellular matrix in cellular processes that ascertain the proper formation of the cell wall in PGs and PTs, but also in other cell types.

## Materials and Methods

### Plant Material and Genotyping

All lines are in the Columbia (Col) background. Seeds were surface-sterilized with 1% sodium hypochlorite, 0.03% TritonX-100, plated and stratified on ½ strength Murashige and Skoog medium (containing 0.6% phytagel, 2% sucrose) for 3 days at 4°C, then transferred to growth chambers with photoperiods of 16 h light and 8 h dark at 22°C. Seedlings were put in pots containing soil and grown in the same growth chamber until flowering. The *LRX* genes share high sequence similarity, which required gene-specific primers (Table S2) for PCR-based genotyping of homozygous mutants. For consistency, only the 5–8^th^ silique on the main inflorescence from at least 12 healthy plants were considered for seed counts.

### Molecular Cloning

All primers used for cloning are listed in Table S2. *LRX8*: two fragments were amplified with the primers pairs LRX8-proF1 + LRX8-terR1 and LRX8-proF2 + LRX8-terR2, digested with *SalI*, ligated and cloned into *pCAMBIA1300* plasmid, which contains kanamycin resistance for selection in bacteria and hygromycin resistance for selection of transgenic plants.

*pLRX11::LRX11*: two fragments named *NT-LRR11* (from the promoter to the end of the cysteine-rich hinge coding sequence) and *EXT11* (from the end of the hinge region to the terminator sequence) were amplified using the primer pairs Lrx11 proF + Lrx11 PstIR and Lrx11 PstIF + Lrx11 terR, respectively. The fragments were ligated, with a *Pst*I site introduced by a silent mutation in the hinge region, into the *pSC* vector (Stratagene) containing an additional *NotI* site introduced at the *XhoI* site to form *pSC-LRX11*. Then the *pSC-LRX11* was cut with NotI, and cloned into the *NotI* site of the binary plant transformation vector *pBART* (Gleave, 1992). *pLRX11::LRX11-Citrine*: the *Citrine* coding sequence (CDS) was amplified with the primer pair T.Cit-F_PstI + Cit-R_PstI, cloned into the PstI site of *pSC-LRX11* to create *pSC-LRX11-Citrine*, and the *LRX11-Citrine* fragment was subcloned into the *NotI* site of *pBART. pLRX11::LRR11-Citrine*: the *LRX11* terminator and the *Citrine* CDS containing a stop codon were PCR amplified with the primer pair PstI 11TF + *NotI* 11TR-1 and T.Cit-F_PstI + T.Cit-R_Pstl, respectively, and cloned into *pSC-pLRX11::LRR11* to create *pSC-pLRX11::LRR11-Citrine*, then subcloned into *pBART. p35S::LRR11-Citrine*: the coding sequence of *LRR11-Citrine* was amplified by PCR from *pLRX11::LRR11-Citrine* with the primers LRR11_oE_F introducing an *XhoI* site 5’ of the ATG start codon and Cit-R_PstI. This fragment was cloned into *pART7* containing a *p35S* promoter and *OCS* terminator (Gleave, 1992) by digestion with *XhoI* and PstI. The resulting overexpression cassette was cloned into *pBART* by *NotI* digestion.

We were unable to clone *LRX9* and *LRX10* due to similar problems we encountered previously with *LRX3–LRX5* (Draeger et al., 2015).

*YC3.60: ProAct1YC3.60* was cut with *HindIII* and *EcoRI* from *pBI121-ProAct1::YC3.60* (Iwano et al., 2009) and cloned into *pSC-LRX11* digested with *HindIII* and *EcoRI* to form *pSN-pAct1::YC3.60*. The *NOS* terminator sequence flanked by *EcoRI* sites was cut and cloned into the *EcoRI* site of *pSN-pAct1::YC3.60*. The *pAct1::YC3.60::NOS* cassette was then subcloned into the *NotI* site of *pBART*.

*nlsR_m_-2A-secG_f_*: the *pSN-pAct1::YC3.60::NOS* was PCR-amplified with the primer pair TNF001 + TNF004 and digested with *BglII* to form a fragment (**a**) lacking the *YC3.60* CDS. The *nlsR_m_-2A-secG_f_* cassette was amplified with the primer pair TNF002 + TNF003, digested with *BamHI*, and ligated to fragment (**a**) to form *pSN-pAct1::nlsR_m_-2A-secG_f_::NOS*. The *pAct1::nlsR_m_-2A-secG_f_::NOS* was then subcloned into the *Not*I site of *pBART*.

The actin-binding GFP construct was kindly provided by Dr. Anna Nestorova (University of Zurich).

Plant transformation and selection of transgenic plants was performed as described previously (Baumberger et al., 2001).

### Quantitative RT-PCR

Open flowers of four independent plants per genotype were used for total RNA extraction by the SV total RNA isolation system kit (Promega) and 300 ng of RNA was reverse transcribed using the iScript advanced cDNA kit (BioRad). qRT-PCR was performed on a CFX96TM real-time system (BioRad) with the Kapa Syber ^®^ Fast qPCR (Kapa Biosystems) technology. *EF α, GAPDH*, and *UBI10* were used as internal standards to quantify expression.

### Alexander Staining

Flowers that had opened on that day were collected and incubated overnight in Alexander staining solution (Alexander, 1969), and cleared for 2 h in chloral hydrate clearing solution containing 8g chloral hydrate, 3 ml glycerine, and 1 ml double distilled water. The anthers were dissected and imaged under DIC using a Leica DMR microscope equipped with a Zeiss Axiocam 105 colour camera.

### Pollen germination and pollen tube growth

Flowers (for good and reproducible germination, mainly the 2 freshly open flowers from the main stems of 4–5½ weeks old plants) from at least 8 plants per genotype were collected and incubated in a moisture chamber for 30 min at 30°C. The liquid PGM (Boavida and McCormick, 2007), pH 7.5, contained 5 mM CaCl_2_, 5 mM KCl, 1.62 mM H3BO3, 1 mM MgSO4, and 10% (w/v) sucrose. Pollen were brushed on silane-coated glass slides (Science Services, <u>www.scienceservices.de</u>) and covered with PGM, germinated, and grown in a moisture chamber at 22°C. To investigate the effect of lower extracellular [Ca^2+^], the concentration of CaCl_2_ in the PGM was reduced to 2 mM. For Ca^2+^ channel inhibition, PGM was supplemented with 5 μM or 15 μM LaCl_3_. For consistency and comparability between experiments, unless explicitly stated otherwise, PT were analysed 5 hrs post germination.

### Immunolabeling of cell wall epitopes

PTs grown for 5 h on silane-coated slides were fixed in PEM buffer (4% paraformaldehyde in 1 M NaOH, 50 mM PIPES, 1 mM EGTA and 5 mM MgSO4, pH 6.9). For enzymatic digest of selected wall components, fixed PTs were rinsed with sodium acetate buffer (pH 5.5) and incubated for 2 h with enzyme solution [5 U/ml solution of xyloglucan-specific xyloglucanase (Megazyme, E-XEGP) prepared in the same acetate buffer] at 37°C. Enzyme-treated and non-treated fixed samples were rinsed 3x with PBS buffer for 5 min each, and blocked with 4% non-fat milk in the same PBS buffer for 1 h or overnight at 4°C. Samples were incubated at RT for 1 h with a 10x dilution of the rat primary antibody (JIM20, LM2, LM19, LM20, LM6, and LM15) in 4% non-fat milk in PBS buffer, then rinsed 3x in the same buffer, and incubated in the dark at RT with 100x dilution of the anti-rat secondary antibody (Sigma, F1763) for 1 h. Controls included non-digested samples and/or omitting the primary antibody. Samples were washed 3x for 10 min each with PBS buffer, and glycerol-based antifade solution (Agar scientific, AGR1320) was added onto the PTs and imaged with a Leica DM6000 microscope.

### Transient gene expression in *Nicotiana benthamiana*

Tobacco infiltration with Agrobacteria (strain GV3103) containing the *pBART-p35S::LRR11-Citrine* construct was performed as described (Bourras et al., 2015).

### Membrane fraction isolation and Western blotting

Seedlings were grown on standard MS medium as described for ten days and homogenized in liquid nitrogen. 100 μL of 1 % SDS was used to extract total protein from 50 mg fresh weight. To extract membrane fractions, a well-established protocol was used (Jasinski et al., 2001): homogenized samples were suspended in 3 volumes of ice-cold extraction buffer [250 mM sorbitol; 50 mM Tris-HCl, 2 mm EDTA; pH 8.0 (HCl); immediately before use add: 5 mM DTT; 0.6 % insoluble PVP; 0.001 M PMSF; 10 μL/mL Protease Inhibitor Cocktail (Sigma P9599)]. The material was first centrifuged at 5,000g and 10,000g for 5 minutes each at 4°C to remove cell debris. The supernatant was then centrifuged at 40,000 rpm for 1 hour at 4°C and the pelleted membrane fraction was resuspended in [5 mM Kh2PO4; 330 mM sucrose; 3 mM KCl; pH 7.8 (KOH); 0.5 % n-Dodecyl-β-D-maltopyranoside]. The samples were used for SDS-PAGE and Western blotting, where the LRR4-Citrine fusion protein and LHC1a were detected with rabbit antibodies (Torrey Pines Biolabs, #TP401 and Agrisera, #AS01005, respectively).

### Aniline blue staining

Fixed PTs were washed three times with 0.1 M phosphate buffer (pH 8.0) and stained directly before microscopy with 0.1% methyl blue (certified for use as aniline blue; Sigma, St. Louis, USA) solution prepared in the 0.1 M phosphate buffer.

### FM4-64 and FM1-43 staining

After about 3 h of germination, PGM was supplemented with 5 μM FM4-64 or 0.16 μM FM1-43 (Molecular Probes™) and PTs were time-lapse imaged.

### Fluorescence quantifications

For quantification, fluorescence was measured as the mean grey value in Fiji (<u>https://fiji.sc/</u>). For immunolabelling, we quantified the fluorescence within 20 μm of the PTs apex, for FM4-64 and FM1-43, the fluorescence within 5 μm of the PT apex and for nlsR_m_-2A-secG_f_, the RFP signal in the nucleus *vs* the GFP signal in the apical cytoplasm.

### Yellow cameleon 3.60 imaging

Time lapse images of PG or PT in PGM inside a humid glass bottom-well petri dish (Mattek) were acquired with an Olympus IX81-ZDC2 inverted wide-field microscope with a CFP/YFP/DsRED filter using a single band excitation filter (436/10 nm) and single band emission filters (465/25 nm and 535/30 nm) at 5 s intervals. Signals were detected by a Hamamatsu EM-CCD camera C-9100, and ratiometric analysis was performed using MATLAB.

### Transmission electron microscopy (TEM)

A detailed step-by-step description of the protocol used was described previously (Ndinyanka Fabrice et al., 2017). Briefly, PT specimens were fixed in 1.25% glutaraldehyde in 0.05% cacodylate buffer, post-fixed in 2% OsO_4_, dehydrated in acetone, and then embedded in Epon. Ultrathin sections as shown in Figure 4C used for measurements of cell wall thickness were collected between 5-15 μm from the PT tip, corresponding to the region where CFM was performed. The sections were visualized in a CM100 TEM system (FEI, The Netherlands) using a Gatan Orius 1000 CCD camera (Gatan, Munich, Germany).

### Cellular force microscopy (CFM)

CFM measurements of apparent stiffness were performed as described (Vogler et al., 2013). Briefly, sensor tips (FemtoTools, Switzerland) were manually positioned on PTs adhering to a silane-coated slide with DIC optics on an Olympus IX 71 inverted microscope (<u>www.olympus-global.com</u>) at the starting point of the measurement series, and then control was taken over by the LabVIEW software. Turgid PTs (n≥28) were indented by a maximum sensor-applied force of 4 μN. At each point, four measurements (with four scans each) were taken from which the mean apparent stiffness value was calculated in MATLAB as previously described (Routier-Kierzkowska et al., 2012).

### Finite element method (FEM) modelling

The extraction of the mechanical properties is performed by fitting the force-indentation curves obtained from the CFM with those acquired from the FEM model. The linear nature of the force-indentation depth curve allows for a single parameter characterized by its slope. The model built for the indentation simulation along with its accompanying uncertainty analysis is identical to that described for other Arabidopsis mutant PTs (manuscript in preparation).

### Accession Numbers

*LRX8*: At3g19020; *LRX9*: At1g49490; *LRX10*: At2g15880; *LRX11*: At4g33970.

## Supplemental Data Files

**Figure S1.** LRX protein structures and effects of mutations on seed set.

**Figure S2.** Protein alignment of LRX8, LRX9, LRX10, LRX11, and LRX4.

**Figure S3.** *In vitro* and semi *in vivo* pollen germination after 5 h incubation.

**Figure S4.** Ultrastructure of pollen grains.

**Figure S5.** Quantification of immunolabeling cell-wall epitopes.

**Figure S6.** Cytoskeletal organization and cytoplasmic accumulation of secretory vesicles.

**Figure S7.** Endocytosis rate in wild-type and *lrx* mutant pollen tubes.

**Figure S8.** Mechanical properties of wild-type and *lrx* mutant pollen tubes.

**Table S1.** Reciprocal crosses.

**Table S2.** List of primers.

**Movie S1:** Bursting of *lrx8/9/11* PGs.

**Movie S2:** Discharge of cytoplasmic membrane-stained components from growing *lrx8/9/11* PT.

**Movie S3:** Vesicle budding form growing *lrx8/9/11* PT.

## Acknowledgments

We are grateful for TEM support from the Centre for Microscopy and Image Analysis of the University of Zürich and thank Prof. Guo (Hebei Normal University, China) for providing the *LRX8* genomic construct, Prof. Clara Sánchez-Rodríguez (ETH Zurich) for the transgenic plant containing the *nlsR_m_-2A-secG_f_* construct, Dr. Anna Nestorova (University of Zurich) for providing the actin-binding GFP construct, and Noemi Peter (University of Zurich) for anti-LHC1a and anti-FBP antibodies (Abcam, Switzerland).

